# Clonal behaviour of myogenic precursor cells throughout the vertebrate lifespan

**DOI:** 10.1101/2022.02.17.480906

**Authors:** Simon M. Hughes, Roberta C. Escaleira, Kees Wanders, Jana Koth, David G. Wilkinson, Qiling Xu

## Abstract

To address questions of stem cell diversity during skeletal myogenesis, a Brainbow-like genetic cell lineage tracing method, dubbed *Musclebow*, was derived by enhancer trapping in zebrafish. It is shown that at least 15 muscle precursor cells (mpcs) seed each somite, where they proliferate but contribute little to muscle growth prior to hatching. Thereafter, dermomyotome-derived mpc clones rapidly expand while some progeny undergo terminal differentiation, leading to stochastic clonal drift. No evidence of cell lineage-based clonal fate diversity was obtained. Neither fibre nor mpc death was observed in uninjured animals. Individual marked muscle fibres persist across much of the lifespan indicating low rates of nuclear turnover. In adulthood, early-marked mpc clones label stable blocks of tissue comprising a significant fraction of either epaxial or hypaxial somite. Fusion of cells from separate early-marked clones occurs in regions of clone overlap. Wounds are regenerated from many/most local mpcs; no evidence for specialised stem mpcs was obtained. In conclusion, our data indicate that most mpcs in muscle tissue contribute to local growth and repair and suggest that cellular turnover is low in the absence of trauma.

**Summary Statement:** Musclebow clonal cell lineage analysis is introduced to reveal the cellular dynamics of skeletal muscle formation, repair and maintenance throughout the life of zebrafish.

## Introduction

Gradual cell lineage restriction is a major determinant of animal development. Understanding cell lineage origins and diversity is also important for designing regenerative stem cell therapies. Clonal analyses have revealed the importance of cell lineage-based behaviours in vertebrates in polarised epithelial tissues such as skin, gut and central nervous system, with stem cell clones occupying a specific niche and contributing to a defined cascade of progeny in a defined planar spatial domain. In contrast, much less is known in mesenchymal tissues with a complex three-dimensional structure. Here, we show how clones of tissue-restricted stem cells contribute stably to patches of skeletal muscle tissue throughout the life of an animal.

Distinct myogenic precursor cell (mpc) lineages have long been proposed to be a major factor controlling vertebrate myogenesis (Cossu et al., 1988; Hauschka, 1974; Hughes and Salinas, 1999; Miller and Stockdale, 1986). Mpc clonal behaviour in cell culture first led to the suggestion that heritably-robust intrinsic differences exist between myoblast clones in embryonic muscles, and that these underpin cell fate decisions, such as the formation of slow and fast fibres (Schafer et al., 1987). Moreover, myoblast populations appear to change character as muscle matures, leading to the suggestion that distinct generations of fibres arise from distinct clones (Hauschka, 1974; Hutcheson et al., 2009; Messina et al., 2010). Clone lineage has also been proposed to control the timing of myoblast differentiation into muscle fibres (Quinn et al., 1985). The essence of these ideas is that of commitment; that a cell (in this case a dividing myoblast) already knows what it will do at the next step in its development. Despite these indications of the importance of cell lineage in vertebrate myogenesis, in vivo evidence is sorely lacking.

Mature vertebrate muscle fibres are multinucleate, formed by the fusion of mononucleate myoblast-derived differentiating myocytes (Collins et al., 2005; Gearhart and Mintz, 1972; Stockdale and Holtzer, 1961). This fact complicates lineage analysis because transplantation and heterokaryon experiments all suggest that upon fusion of a myocyte to a pre-existing fibre, the myocyte nucleus is reprogrammed to copy the host fibre gene expression (Blau et al., 1985). Moreover, clonal analyses in vivo demonstrate that most amniote mpcs fuse randomly with fibres in their environment (Hughes, 1999; Hughes and Blau, 1990; Hughes and Blau, 1992; Robson and Hughes, 1999). More recently, renewed interest in the lineage hypothesis has arisen from two observations. First, it is clear that muscle satellite cells, the tissue-restricted stem cells of muscle growth and repair in adult animals, are derived from the somite and share some, but not all, of their molecular characteristics with early myoblasts (Cossu et al., 1988; Gros et al., 2005; Kassar-Duchossoy et al., 2005). Secondly, muscle shows a remarkable robustness to genetic manipulations, suggesting that extrinsic signals can, in appropriate circumstances, override the intrinsically-encoded destiny of a myoblast (Haldar et al., 2008). Although distinct populations of differentiated muscle fibres have been suggested to be formed from committed but proliferative mpc sub-populations, evidence from unmanipulated endogenous mpcs is not compelling (DiMario et al., 1993; DiMario and Stockdale, 1995; Hughes, 1999; Hughes and Blau, 1992; Motohashi et al., 2019). Recent efforts have employed Zebrabow technology to analyse cell lineage during zebrafish muscle development and regeneration (Gurevich et al., 2016; Nguyen et al., 2017). These studies have shown that 1) a few clonally-related myoblasts contribute to muscle regeneration in larval somites (Gurevich et al., 2016) and 2) neutral clonal drift sometimes causes single stem cells to label large portions of the myotome as animals grow (Nguyen et al., 2017). Zebrabow constructs have multiple insertions and express in all cell types, which can make tracing individual clones difficult (Nguyen and Currie, 2018). In order to test hypotheses implicating cell lineage as a regulator of muscle development, we developed an alternative approach in zebrafish that enabled analysis of mpc behaviour. We carried out an enhancer trap screen using the *Brainbow-1.0 ‘L’* construct (Livet et al., 2007) and found a line in which expression is restricted to somitic muscle lineages, which we term Musclebow. Recombination to generate different fluorescent colours was achieved by short heat shock induction of Cre recombinase expression.

Here we describe Musclebow lineage analysis of somite cells throughout the zebrafish lifespan. Using this approach, we tracked large numbers of single muscle fibres and somite cell clones over hours, weeks and years. We found that myocytes are initially highly dynamic during their differentiation from muscle stem cells, but then become stably incorporated into myotomal structure. Marked clones of motile mononucleate somitic cells disperse through the somite from the dermomyotome, following stereotypical routes. Clones marked at embryonic stages are generally contained within single somites and contribute to regions of muscle constituting a broad zone of fibres. There is significant clonal drift over time, leading to large clones restricted to single contiguous regions of individual somites that appear stable throughout adult life.

## Materials and Methods

### Zebrafish husbandry

Transgenic lines were created on London/AB wild type background and backcrossed onto AB. Maintenance, staging and husbandry were performed as previously described (Westerfield, 2000). Lines used are listed in Table 1.

**Table 1.**
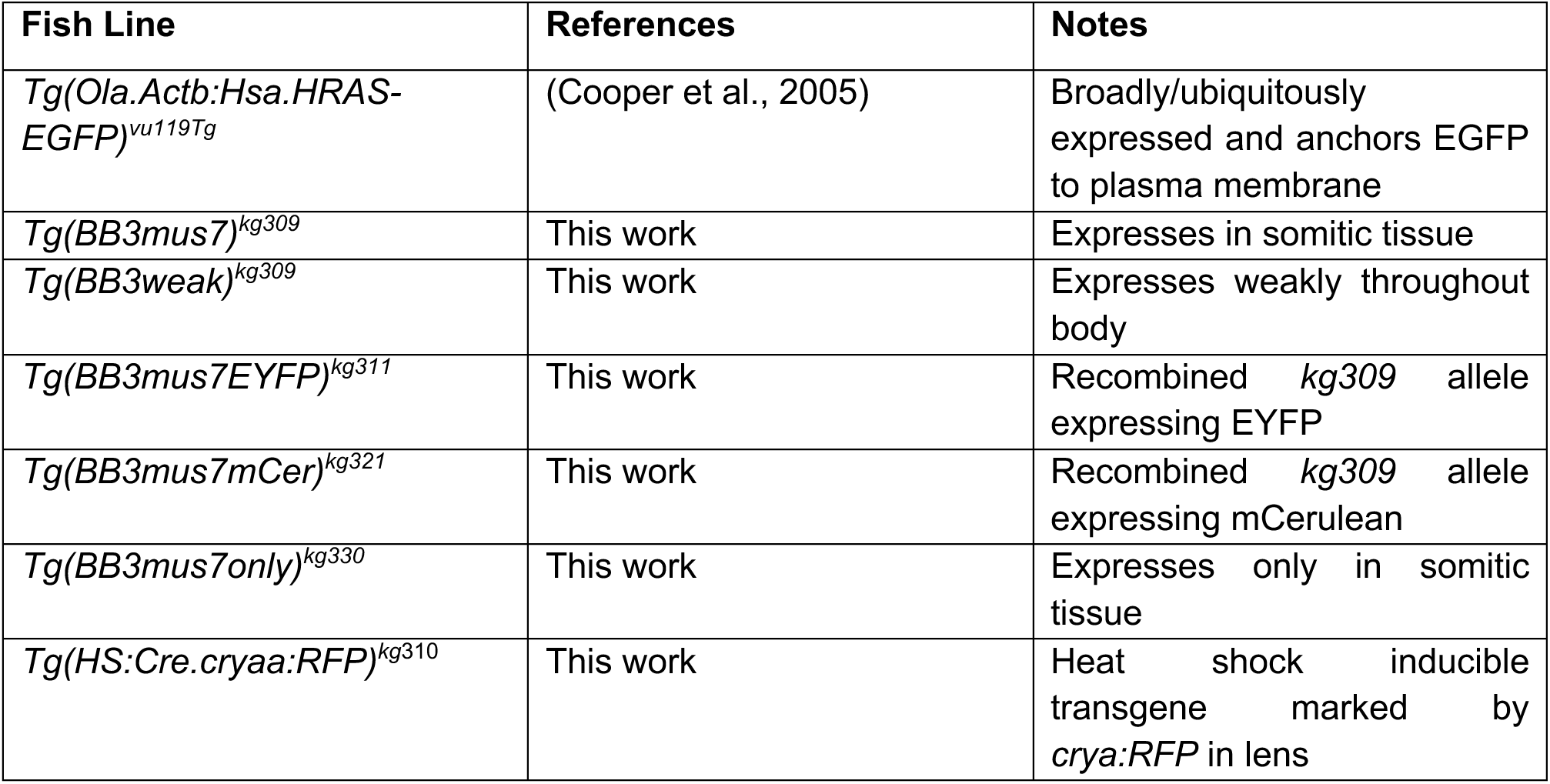
Fish alleles.

### Zebrafish Transgenesis

Transgenic Musclebow zebrafish lines were made with a Tol2-based enhancer trap construct encoding three fluorescent proteins (tdTomato, mCerulean and EYFP) flanked by paired loxP and loxP2272 sites, under the control of a 1.2 kb *claudin-b* basal promotor (Fig. 1A), based on the *Brainbow-1.0 ’L’* vector (Livet et al., 2007; Pan et al., 2013). One line generated, designated *Tg(BB3mus7)^kg309^*, had strong fluorescent signal in muscle fibres and weak signal elsewhere, each of which showed linked monogenic Mendelian transmission, and was analysed further. The tdTomato detected diminished with age of the fish, perhaps due to an integration site effect and/or overall reduction in transcription/translation rate in mature muscle fibres. Upon injection of RNA encoding Cre into *Tg(BB3mus7)^kg309^* embryos, both EYFP and mCerulean (mCer) marked cells were clearly visible, although the former were more abundant. mCer-marked cells generally appeared dimmer than EYFP-marked cells. A rare recombination event during outcrossing of *Tg(BB3mus7)^kg309^* to wild type created a subline with tdTomato expression more restricted to somites and muscle, and was designated *Tg(BB3mus7only)^kg330^* (Fig. S1). In general, *Tg(BB3mus7)^kg309^* larvae lacking the linked *Tg(BB3weak)* insertion present in the parental *kg309* allele were selected for experiments.

**Fig. 1.**
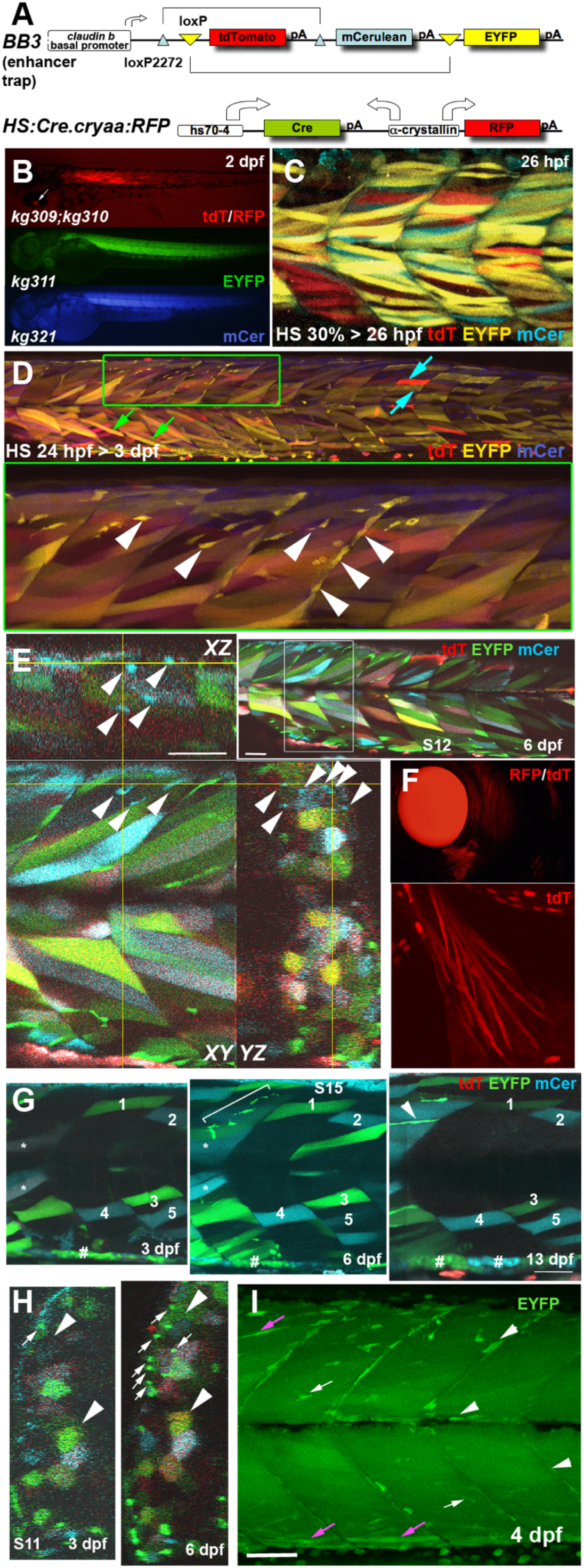
Musclebow fish allow genetic marking of muscle fibres and their precursor cells. Transgenic Musclebow fish were selected from an enhancer trap screen with larvae at the indicated stages shown in lateral views with anterior to left, dorsal up. **A.** The BB3 and HS:Cre.cryaa:RFP DNA constructs used for making transgenic fish. **B.** Dissecting scope view of *BB3mus7^kg309^* yielding tdTomato detectable in a subset of muscle fibres from 1 dpf and *HS:Cre.cryaa:RFP^kg310^* yields RFP in lens from 2 dpf (arrow, top). Germline activity of *HS:Cre.cryaa:RFP^kg310^* yielded lines *BB3mus7:EYFP^kg31^*^1^ (EYFP, middle) or *BB3mus7:mCerulean^kg321^* (mCer, bottom) expressing throughout muscle tissue. **C.** Maximum intensity projection confocal stack (mip) after heat shock at 30% epiboly yielding subsets of fibres expressing one or two recombined and unrecombined alleles, presumably reflecting mosaic Cre action followed by fusion of mpcs expressing *BB3mus7* in different recombination states. Note that recombination yielded EYFP fibres more frequently than mCer fibres and that some fibres remain unlabelled. **D.** Three channel confocal mip stack showing *BB3mus7* expression at 3 dpf in morphological slow (cyan arrows) and fast (green arrows) fibres and in mncs (arrowheads) in somites. Note the clustered mncs (green box, short stack shown magnified beneath). **E.** Single cluster of mCer-marked mncs (arrowheads) in somite 9. Yellow crosshairs indicate planes in *XY*, *YZ* and *XZ* projections. Inset shows location of cluster (white box). Note absence of mCer mncs in rest of plane. **F.** *BB3mus7* expression in pectoral fin muscles. **G.** Time-lapse analysis of short 20 μm stack mips showing the persistent set of marked fibres (numbered) in the deep region of somites 15 and 16 at 3, 6 and 13 dpf. The 6 dpf stack extends further superficially within the myotome than the 3 and 13 dpf stacks. A cluster of mncs (bracket) appears in the deep myotome between 3 and 6 dpf and forms a nascent fibre (arrowhead) by 13 dpf. Note the loss of tdT in dual labelled fibres (asterisks) and the clonal growth of gut-associated cells (hashes). **H.** Time-lapse analysis of somite 11 shows maintenance of overall pattern of fibre labelling but changes in colour of some fibres (arrowheads). Note dramatic increase in EYFP-marked superficial mncs (arrows) in this somite which correlates with accumulation of extra EYFP-marked fibres (upper arrowheads). **I.** Confocal maximum intensity projection of 4 dpf *BB3mus7:EYFP^kg311^* larva with strong near-uniform signal in myotomal fibres and stronger signal in some mncs on myosepta (arrows) or within the myotome (arrowheads). Note the strongly-marked nascent fibres (magenta arrows). Bars = 50 μm.

Separate Cre driver lines were generated using a construct in which the heat-shock-dependent *hs70-4* promoter drives Cre (Hans et al., 2011) and the *α*-crystallin *cryaa* promoter drives RFP in the lens, permitting live identification of fish carrying the transgene from 2 dpf onwards (Fig. 1A). The *Tg(HS:Cre.cryaa:RFP)^kg^*^310^ driver line was selected and shown to give Cre expression for around 1-3 h after heatshock at 39°C. The *Tg(HS:Cre.cryaa:RFP)^kg^*^310^ driver was bred onto *Tg(BB3mus7)^kg309^*, in which it showed good recombination in a wide range of tissues after early heatshock (Fig. 1). Brief heatshocks of under 5 minutes gave relatively little recombination, permitting clonal analysis of cell fate. Expression of EYFP or mCer was observed in patches of muscle fibres. Transmission of *Tg(HS:Cre.cryaa:RFP)^kg^*^310^ from the mother without heatshock yielded germline recombinants expressing either EYFP (*kg311*) or mCer (*kg321*) widely in skeletal muscle, presumably due to Cre expression during chromosome restructuring in the oocyte (Figs 1B and S1). Therefore, in clonal analysis experiments, *Tg(HS:Cre.cryaa:RFP)^kg^*^310^ was always delivered from the male parent.

### Somite Cell Analysis

RNA encoding Histone2B-mCherry (Shaner et al., 2004) was injected into *Tg(Ola.Actb:Hsa.HRAS-EGFP)^vu119Tg^* at the early one cell stage. To ensure that all nuclei were labelled in the analysed larvae, after scanning each fish was lightly fixed in 4% paraformaldehyde and stained for Hoechst 33342 and re-scanned; although intensity varied, all blue nuclei were also red in the quantified larvae. Quantification was performed in Volocity by marking every nucleus and manually attributing each to either a fibre or mononuclear somitic cell (mnc) in the confocal stack. The presence of EGFP^+^ T-tubule striations defined fibres. Point spread function of light in the *Z*-axis led to weaker but distinguishable plasma membrane signal when imaged *en face*.

### Imaging and Data Processing

Wholemount images were acquired with an Olympus DP70 camera on a Leica MZ16F dissecting microscope and confocal images were acquired in ZEN on a Zeiss LSM Exciter with a 40x/1.1 W Corr LD C-Apochromat objective for fixed specimens. For live time-lapse imaging with a Zeiss 20x/1.0 W Plan-Apochromat, fluorescent embryos were mounted in 0.8-1% low melting point agarose (LMPA) in embryo medium (EM) containing 160 mg/L tricaine as an anaesthetic (Westerfield, 2000) in a Petri dish and covered with EM. *Z*-spacing was generally set to optimal for 1 Airy unit. Tiff stacks were exported to Volocity 4.2-6.3 (Perkin Elmer) for further analysis and are shown as maximum intensity projections unless otherwise stated. EYFP and mCer persisted well during repeated scanning, whereas tdTomato faded faster.

### Muscle Regeneration Assay

Lesions were produced and regeneration monitored as described (Pipalia et al., 2016). Tg(*BB3mus7*)*^kg^*^309^*;Tg(HS;Cre.cryaa:RFP)^kg^*^310^ larvae were embedded in 1% LMPA, imaged under Zeiss confocal, injured by needle-stick in selected epaxial somites and re-scanned at 1-2 hours post wound (hpw), released from LMPA and kept in a standard incubator at 28.5°C. At various time-points from 1-4 days post wound (dpw) injured larvae were re-embedded and scanned to trace the behaviour of marked cells during wound repair, being released from LMPA at least once every 24 h.

## Results

### Musclebow cell lineage tracing

Musclebow transgenic fish were generated with Tol2-mediated enhancer trap (Fig. 1A). Transgene insertion of the BB3 construct led to *Tg(BB3mus7)^kg309^*, which initially expressed tdTomato (tdT) in what appeared to be a random subset of muscle fibres and mononucleate somitic cells (mncs, which in our definition do not include mononucleate muscle fibres) in larval fish (Fig. 1B). The *BB3mus7* transgene could undergo germline recombination to express either EYFP or mCer in the presence of a second transgene (*Tg(HS:Cre.cryaa:RFP)^kg310^*) expressing Cre recombinase and labelled the entire myotome (Fig. 1B). Later Cre expression led to multi-coloured muscle fibres (Fig. 1C). Importantly, mononucleate superficial slow fibres and mncs never contained both EYFP and mCer, indicating that only a single BB3 transgene was present at the *Tg(BB3mus7)^kg309^* enhancer-trap locus. The presence of fast fibres with a variety of colour combinations, including both EYFP and mCer, therefore indicates the fusion into single multinucleate fibres of nuclei expressing distinct colours (Fig. 1C-E). Expression was stable over time (Movie S1). Many marked mncs (either EYFP- or mCer-marked) appeared to be mpcs based on their morphology and location (Fig. 1D; (Nguyen et al., 2017; Roy et al., 2017)). Marked mncs were arranged in clusters, suggesting a clonal origin, and tended to be brighter than neighbouring marked fibres, presumably due to their lower cytoplasmic volume concentrating the EYFP (Fig. 1D). Early-marked mnc clusters appeared to span one or several adjacent somites, but tended to be restricted to either epaxial or hypaxial domains (Fig. 1D). Mnc clusters labelled with mCer were rarer than those marked with EYFP, but mnc clusters of each colour were generally spatially well-separated (Fig. 1E). For example, examination of an entire confocal stack spanning somites 7 to 17 (S7-17) revealed only three ‘clusters’ of mCer-marked mncs, one of 13 cells restricted to the epaxial regions of S8+S9 (Fig. 1E), a second of 4 cells in the hypaxial extremes of S7+S8, and a third of 5 cells restricted to the hypaxial region of S12 (data not shown). The chance of this clustering of 22 marked cells into ≤5 out of 20 somitic halves is <1/10^6^, strongly supporting a clonal origin of most cells in clusters. From static images, however, we cannot be certain that any individual cluster is clonal (two cells near to one another could have been converted to mCer independently), as we have shown in previous muscle lineage-tracing studies. Nevertheless, when clusters are well-spaced and of a single colour, behaviours of clusters that are often observed are highly likely to arise within single clones (Hughes and Blau, 1990; Hughes and Blau, 1992). Mnc clusters (either mCer- or EYFP-marked) were less frequent in more posterior somites, perhaps reflecting the graded decreases in somite size and cell number along the body axis. Mnc clusters were rare compared to marked fibres, a finding to which we return below.

*Tg(BB3mus7)^kg309^;Tg(HS:Cre.cryaa:RFP)^kg310^* fish were repeatedly analysed over periods of days and weeks through capture of 3D confocal stacks of a defined group of 3-4 somites, or 3D tile scanning of entire 31 somite blocks. Single marked muscle fibres identified at 3 dpf could be tracked into later stages with great confidence (Fig. 1G). Individual fibres identified in a specific region of an early somite generally retained their shape, orientation and fluorescent protein expression at later stages. No cases of muscle fibre death were observed in many (>15) larvae examined by time-lapse, showing that fibre death is rare. Occasionally, identified fibres changed colour somewhat, relative to surrounding cells, presumably reflecting fusion of mpcs of a distinct colour to the tracked fibre (Fig. 1H). Labelled fibres were also observed in head and pectoral fin muscles (Fig. 1F). Germline recombined *Tg(BB3mus7EYFP)^kg311^* showed many intensely-marked mncs despite strong and rather uniform labelling in fibres (Fig. 1I). Marked mncs were in the typical locations of mpcs near horizontal and vertical myosepta, between fibres within the myotome and appeared to be forming nascent fibres at 5 dpf (Fig. 1I). We conclude that *Tg(BB3mus7)^kg309^* permits tracking of mpc proliferation and differentiation during growth of zebrafish skeletal muscle.

### Cell content of the larval somite

To interpret clonal analyses quantitatively, it is advantageous to know the numbers of mncs, fibres and their nuclei within the somite. We therefore injected RNA encoding Histone2B-mCherry into *Tg(ßactin:Hras-EGFP)^vu119Tg^* fish to label nuclei red and plasma membrane green and analysed the cells within the somite at 3 dpf in high-resolution confocal stacks (Fig. 2). Fibres were readily distinguished from mncs by position and the presence of T-tubule striations (Fig. 2; Table 2). At this stage, 31 total mononucleate fibres were present in somite 16, of which around 23 were the superficial slow fibres. An additional 64 multinucleate fibres with an average of 3.0 nuclei/fibre gave a total fibre content of 96 fibres (Table 2). Fast fibres had between one and six nuclei, with a mean of 2.8 nuclei/fibre and a median and mode of 3 nuclei/fibre (Table 2). Mncs in somite 16 numbered 89, 28% of total somitic nuclei. At 3 dpf, no mncs were detected within the body of the myotome; most mncs were in the dermomyotomal layer (60), particularly at the dorsal and ventral somite extremes, or at the vertical (17) or horizontal (12) myosepta (Fig. 2; Table 2). Epaxial and hypaxial regions of the somite had very similar cellular composition (Table 2). However, the hypaxial domain had slightly more and larger muscle fibres and slightly fewer mncs (Fig. 2; Table 2). While every effort was made to distinguish epi/peridermal cells from cells at the lateral somite surface, we cannot be confident that cells such a neural crest-derived xanthophores, located between somite and epidermis, were always excluded, as their flat morphology is similar to that of dermomyotome cells; such inclusion might contribute to the larger numbers of mncs observed in the epaxial domain. Based on analyses of fluorescently-marked xanthophores in somite 16 at this stage (Mahalwar et al., 2014; Pipalia et al., 2016), there might be up to six in epaxial and four in hypaxial domain. Analysis of other 3 dpf fish showed similar cell and nuclear numbers but varied between and within lays in parallel with the absolute size of the larvae and thus of somite 16 (Kelu et al., 2020). Thus, at 3 dpf, over two thirds of somitic nuclei are in terminally-differentiated muscle fibres, but around a quarter of all nuclei are in potentially-myogenic mncs.

**Fig. 2.**
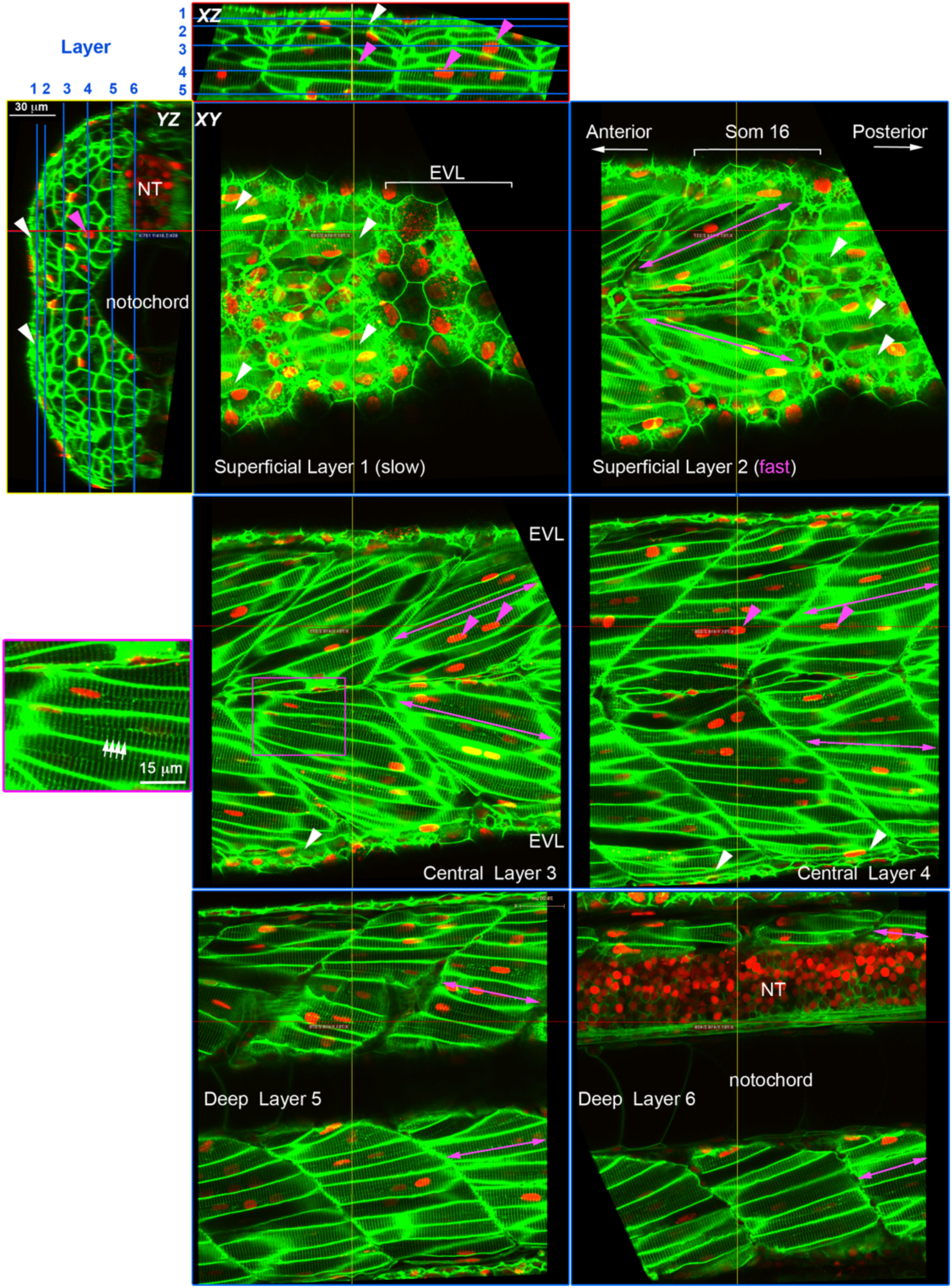
Cell content of the growing zebrafish somite. Larvae expressing membrane-localized EGFP and injected with RNA encoding H2B-mCherry were selected for brightness of both labels and scanned live at high resolution using a sub-Airy pinhole. Three orthogonal views are shown, with blue lines on *XZ* and *YZ* projections indicating the six sagittal *XY* planes. Red and yellow lines indicate the XZ and YZ plane, respectively. Note that EGFP labels the transverse T-tubule system (white arrows) in muscle fibres at regular 2 μm spacing orthogonal to the long axis of the fibre (purple box, magnified at left). Mononucleated fibres parallel to the anteroposterior axis on the surface of the myotome (white arrowheads) are superficial slow fibres. Fast fibres with multiple nuclei (magenta arrowheads) show different orientation depending on the depth within the myotome (magenta arrows); superficially they ‘point’ anteriorly, but in the medial myotome they ‘point’ posteriorly. NT neural tube.

**Table 2.** Cellular content of Somite 16 at 3 dpf.

| Somite 16 | Fraction of nuclei (%) | Number of Cells (number of nuclei) |  |  |
| --- | --- | --- | --- | --- |
|  |  | Epaxial | Hypaxial | Total |
| Slow mononucleate fibres | 7 | 10 (10) | 13 (13) | 23 (23) |
| Fast mononucleate fibres | 3 | 3 (3) | 5 (5) | 8 (8) |
| Fast binucleate fibres | 13 | 9 (18) | 12 (24) | 21 (42) |
| Fast trinucleate fibres | 23 | 11 (33) | 13 (39) | 24 (72) |
| Fast tetranucleate fibres | 22 | 8 (32) | 9 (36) | 17 (68) |
| Fast penta/hexanucleate fibres | 4 | 0 (0) | 2 (11) | 2 (11) |
| Fast fibres total | 64 | 31 (86) | 41 (115) | 72 (201) |
| **Total fibres (myotome nuclei)** | **72** | **41 (96)** | **54 (128)** | **95 (224)** |
| Nuclei/fast fibre |  | 2.8 | 2.9 | 2.8 |
| Nuclei/slow fibre |  | 1.0 | 1.0 | 1.0 |
| Nuclei/fibre |  | 2.3 | 2.4 | 2.4 |
| Dermomyotome cells | 19 | 36 (36) | 24 (24) | 60 (60) |
| Vertical myoseptum cells* | 5 | 16/2=8 (8) | 18/2=9 (9) | 34/2=17 (17) |
| Horizontal myoseptum cells | 4 | - | - | 12 (12) |
| **Total non-fibre cells (nuclei)** | **28** | **44 (44)** | **33 (33)** | **89 (89)** |
| **Somite cells (somite nuclei)** | **100** | **85 (140)** | **87 (161)** | **184 (313)** |

### Clonal cell lineage analysis in Musclebow fish

Previous analyses in 24 hpf somites have revealed ∼40-50 Pax3/7-marked mpcs in each somite 12-16 (Hammond et al., 2007; Hollway et al., 2007; Seger et al., 2011; Windner et al., 2012), suggesting significant proliferation between 1 and 3 dpf if most mncs at 3 dpf are mpcs. Congruently, many mpcs are in the cell cycle over this period (Hammond et al., 2007; Roy et al., 2017; Seger et al., 2011; Stellabotte et al., 2007). To analyse the behaviour of mnc clones, cells were marked within early somites and their fates tracked (Fig. 3). Applying a brief heat-shock at or after 24 hpf led to many single labelled muscle fibres (Fig. 3A). Importantly, these included the mononucleate superficial slow fibres (SSFs) that can be identified by their superficial location, orientation parallel to the anteroposterior axis and large central nucleus and muscle pioneer slow fibres (MP) located at the horizontal myoseptum (Fig. 3B). SSFs and MPs are already terminally differentiated at the time of heat-shock (Devoto et al., 1996), indicating that Cre is active inside differentiated muscle cells. Consistent with this view, no discernible clustering of fibres of the same colour was observed shortly after heat-shock (Fig. 3B). Quantification revealed ∼1.6 (range 0-4) marked SSFs and ∼0.2 (range 0-1) marked MP per somite at 3 dpf (Fig. 3C) indicating that <10% of slow fibre nuclei underwent recombination (Table 2). Recombination events in individual somites appeared to be randomly distributed, both in terms of fibre number and whether EYFP or mCer was expressed. A tendency to fewer recombined slow fibres in more caudal somites likely reflected the lower numbers of slow fibres as the trunk tapers into the tail (Fig. 3C). Thus, marked slow fibres represent separate recombination events.

**Fig. 3.**
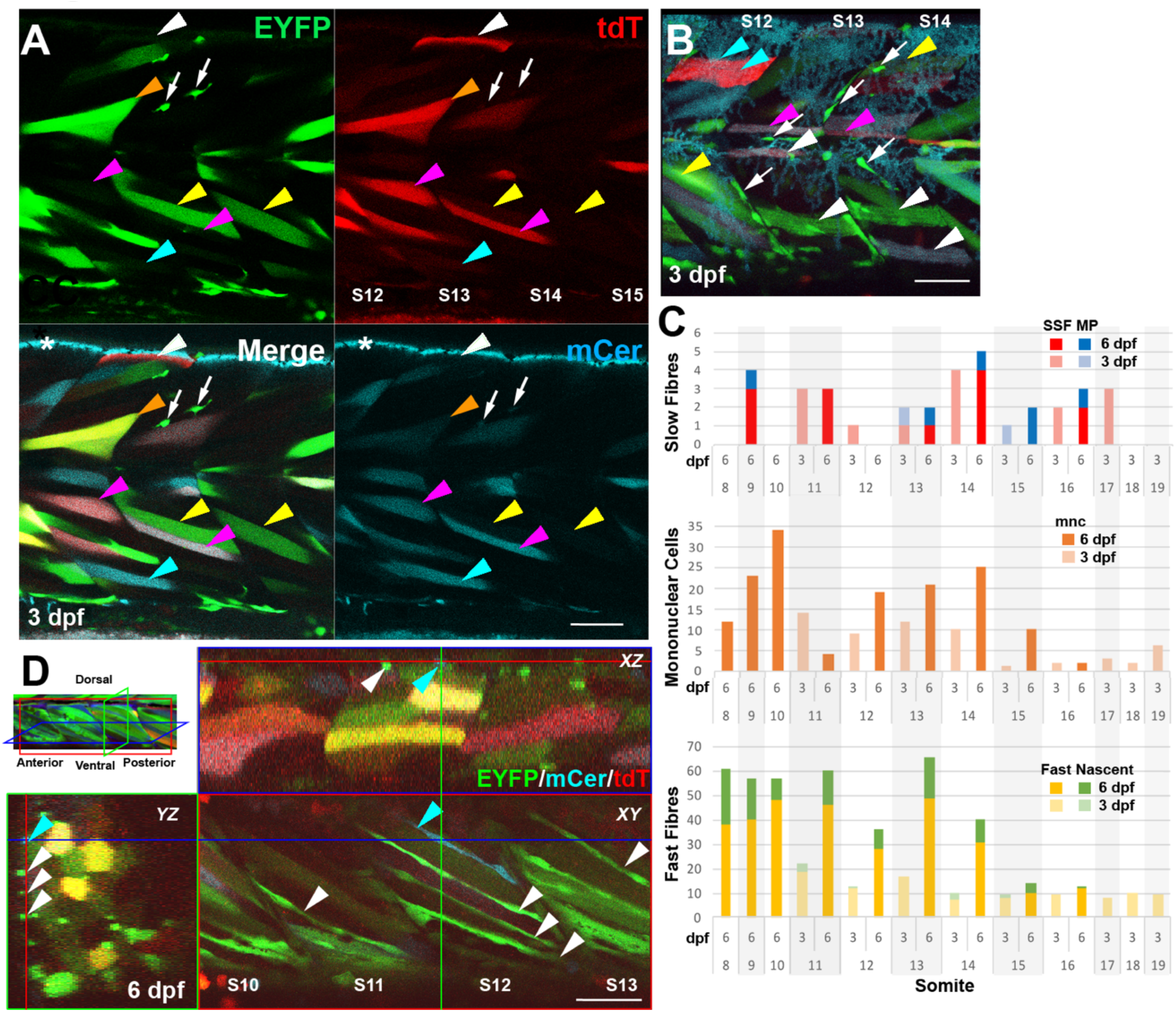
*BB3mus7* recombination reveals dynamics of mpc clones. Live confocal microscopy of *BB3mus7^kg309/+^; HS:Cre.cryaa:RFP^kg310/+^* embryos after heat-shock at 24 hpf. **A.** Three channel confocal slice showing *BB3mus7* expression in somites 12-15 at 3 dpf in morphological slow (white arrowheads) and fast (other arrowheads) fibres and in mncs (white arrows) in somites. Note the clustered mncs. Examples of individual fibres marked by tdT-only (white arrowheads), EYFP-only (yellow arrowheads), mCer-only (cyan arrowheads), EFYP and tdT (orange arrowheads) and mCer and tdT (magenta arrowheads). Asterisk indicates autofluorescence from xanthophores in the mCer channel. Faint breakthrough of bright EYFP signal into the mCer detection channel that can be seen, for example, in the mncs and the fibre indicated by the orange arrowhead is readily distinguished from genuine mCer. **B.** Short stack (22 μm) mip of somites 12-14 at 3 dpf showing recombined mncs (arrows), superficial slow fibres (white arrowheads), muscle pioneers (magenta arrowheads), fast fibres (yellow arrowheads) and unrecombined slow fibres (cyan arrowheads). Note that some fibres contain both tdT and mCer or EYFP proteins, indicating perdurance of tdT after Cre-driven recombination. **C.** Quantification of numbers of marked cells in individual somites of a single fish at 3 and 6 dpf, determined from high-resolution confocal stacks. Only somites 11-16 were scored at both timepoints. SSF, superficial slow fibre; MP, muscle pioneer slow fibre; mnc, mononuclear non-fibre cell; Fast, fast fibre with T-tubules; Nascent, thin nascent fibre spanning anteroposterior extent of somite. **D.** *XYZ* slice projections (indicated on mip at top left) of somites 10-13 at 6 dpf showing reduction in mncs and increase in elongated cells spanning the myotome marked with EYFP (white arrowheads) or mCer (blue arrowheads) within the hypaxial myotome. Coloured crosshairs indicate the plane of the orthogonal boxed optical sections. Bars = 50 μm.

Comparing numbers of recombined slow and fast fibres at 3 dpf showed that marked fast fibres were more numerous than slow, roughly in proportion to the ratios of their nuclei present at the time of heat-shock (Fig. 3A,C; Table 2). Within a larva, the numbers of marked slow and fast fibres were well-correlated, although separate larvae showed different recombination rates. Thus, within one animal, the chance of recombination appears similar for each nucleus. As muscle fibre nuclei are post-mitotic and unable to proliferate further, such marked fibres constitute a reference background pattern facilitating tracking of other marked clones.

In contrast to fibres, clusters of similarly-marked mncs were evident at 3 dpf (Fig. 3A,B) and over ensuing days contributed to many narrow elongated fibres, particularly in the hypaxial somitic extreme (Fig. 3D). The approximately 28% of somite nuclei in mncs at 3 dpf (Table 2) provide a significant pool for later muscle growth, and beg the question of their clonal origin over the preceding days. At one extreme, all somite mpcs might derive from a single original somite cell. At the other, all mpcs might have an independent origin as the somite formed. Marked mncs were rare in some somites and more abundant in others two days after heat-shock, averaging at 6.6 marked mncs/somite (Fig. 3C). This probably reflects stochastic recombination in the low numbers of cells in a somite that have not formed muscle fibres at 24 hpf (Fig. 4A; (Devoto et al., 2006; Hammond et al., 2007; Hollway et al., 2007; Stellabotte et al., 2007)). Moreover, analysis of the distribution of marked mncs across a population of embryos suggested all somites contained mncs that could, on occasion, undergo recombination. Considering the 89 mncs counted in somite 16 (Table 2), we deduce that around 7% (6.6/89) of mncs are marked, a recombinant fraction similar to that of slow fibres and proving that mncs within single somites derive from multiple clones. As the number of marked mncs in individual somites ranged from one to 14, it appears that either different numbers of mncs underwent recombination in each somite and/or individual marked mnc precursors show a range of proliferative behaviours prior to 3 dpf. Strong support for the latter view comes from the rarity and scattered distribution of mCer-marked mnc clusters, which nevertheless also contain 2-8 cells/somite, as mentioned above. The similar cluster size, despite threefold lower overall frequency argues that many mnc clusters are clones.

**Fig. 4.**
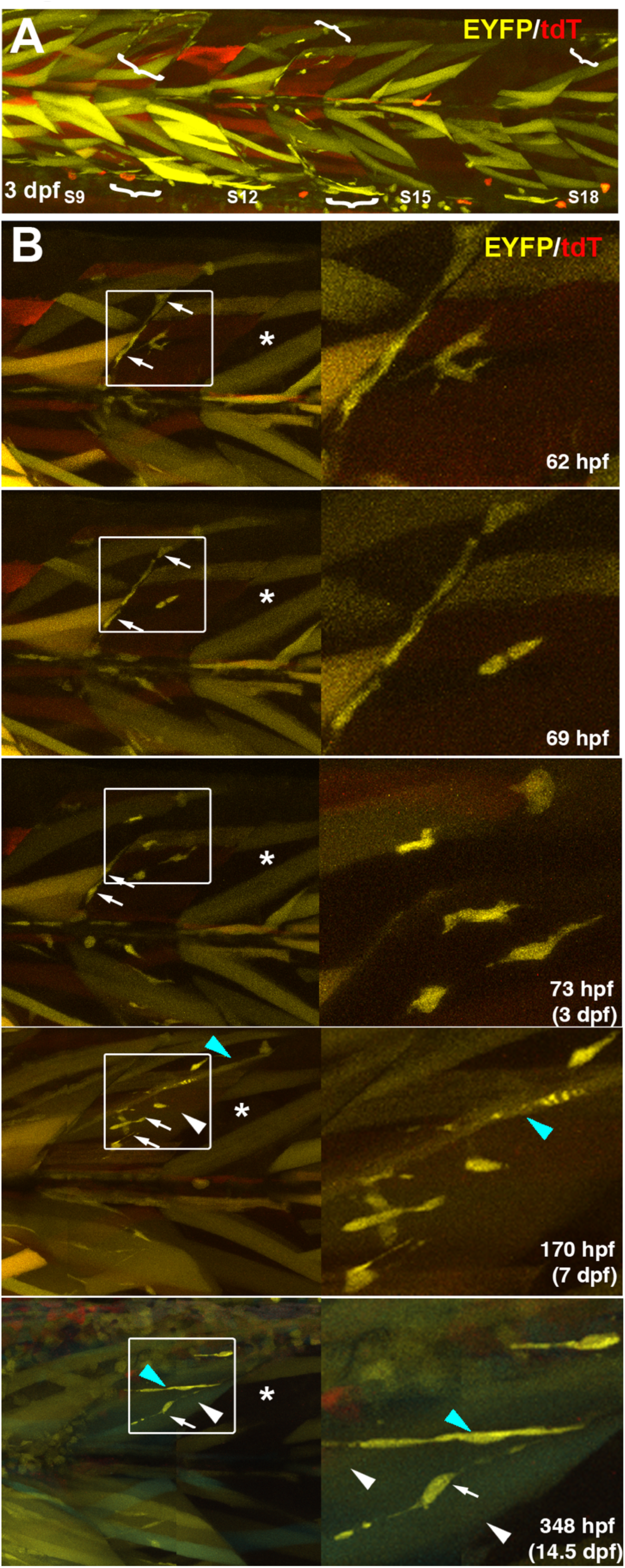
Kinetics of proliferation and differentiation of an EYFP-marked mnc cluster. Time-lapse series taken from full somite three channel confocal stacks of a single Musclebow marked fish after heat-shock at 24 hpf. **A.** Full somite stack mip of tile scan of somites 9-18 showing five clusters of EYFP-marked mncs (brackets) at 3 dpf. Note the large areas lacking marked mncs and the more numerous marked fibres in anterior somites. **B.** Time-lapse confocal mips showing development of the mnc cluster in somite 14 of panel A. Boxes shown magnified at right. Note the i) early presence and later absence of EYFP-marked mncs on the vertical myoseptum (arrows), ii) increase in number of marked mncs, nascent fibres (blue arrowheads) and large fibres (white arrowheads) in the body of the myotome and iii) lack of increase in marked fibres in the adjacent region of somite 15 that also lacked marked mncs (asterisks).

Although there was a trend towards higher marked mnc numbers in the larger more rostral somites. mnc labelling was still rare (Fig. 3C). Clusters of marked mncs were often restricted to single half somites (epaxial or hypaxial) and did not appear to cross the vertical myoseptum into adjacent somites (Fig. 4A), points to which we return below. Based on the frequency of fibre recombination and that of marked mnc clusters, we conclude that a pool of at least 15 cells generate the mncs in each trunk somite 8-19.

Repeated imaging of individual fish revealed that clusters of marked mncs persisted in specific regions within individual somites as fish matured, but showed varied behaviour. Such mnc clusters usually had a single colour, consistent with clonal origin (Figs 3A,B,D). Although clusters were rare in fish showing low recombination levels, over time clusters grew larger, consistent with proliferation of clones (Fig. 3C). Indeed, on occasion, short time-lapse movies captured cytokinesis of mncs within a cluster (Fig. 4B). Quantification showed that mnc numbers rose in somites 11-16 by 70% on average between 3 and 6 dpf, although outcomes were heterogeneous. Marked mnc number in somite 11 dropped from 14 to 4, whereas in somite 15 it rose from 1 to 10 (Fig. 3C). A lack of change in mnc number was rare. Thus, the behaviour of marked mncs is dynamic with apparently stochastic increases and decreases in cluster size.

Analysis of the distribution of mnc clusters suggested that the vast majority represent clones. As already mentioned, clusters were generally homogeneous in colour. For example, in one fish at 3 dpf, only two out of nine somites analysed contained mCer-marked mncs; somite 17 had a cluster of three mncs in the ventral region but lacked EYFP-marked mncs and somite 18 had one mCer-marked mnc in the hypaxial dermomyotome and one EYFP-marked mnc at the dorsal edge of the epaxial region. The chance of obtaining clustering of at least 3 out of 4 cells in the same half-somite if each mCer^+^ mnc arose from an independent recombination is 0.012. However, mCer mnc clusters were, on average, both fainter and smaller than EYFP clusters, suggesting that the weaker blue label prevented detection of all cells in a clone; we therefore focussed on EYFP clusters. In one embryo, for example, in which the dorsal half of all 31 somites was imaged at high resolution, only a single somite had marked mncs. Yet, this epaxial somite had at least four marked mncs. Assuming an equal number of target cells in each somite, the chance of each of these cells being a separate recombination event and yet being located in the same somite is (1/31)^4-1^ i.e. p < 4 x 10^-5^. Even the presence of two clones in this half somite is unlikely (*p* = 0.032). Similar non-random distributions were observed in many embryos. Thus, we can be confident that, so long as overall numbers of marked mncs are low, most clusters are clones.

In summary, the data indicate that:

a. fewer than 10% of all mncs are marked within the analysed Musclebow larvae. This means that Cre recombination frequency is low – most mnc precursors did not become labelled (because brief heat shock activated low levels of Cre in just a few cells).
b. once recombined and expressing EYFP, mncs tend to retain expression over a significant period. Given that marked fibres remain stably labelled for weeks, we have no evidence for reporter shutdown once a cell is recombined and expressing the transgene at detectable levels.
c. mncs in somites arise from multiple original somite cells. When heat shock was applied shortly after somite formation, only a fraction of mncs in a subset of somites were labelled. This result indicates that many mncs are not derived from the marked mnc clones, and is consistent with the described abundance of Pax3/7-marked cells in somites at 24 hpf (Hammond et al., 2007; Stellabotte et al., 2007).
d. most somite mncs undergo divisions by 3 dpf. To label a small clone within a somite, several cell divisions of the original marked cell must have occurred in the two day period. As few somites contain single marked mncs (as would be expected if mncs became marked but did not divide), most mncs seem to divide between 1 and 3 dpf.
e. mncs rarely form new fibres between 1 and 3 dpf. The absence of marked elongated fibres next to mnc clones suggests few cells form new fibres. Fusion to large pre-existing fibres cannot be ruled out because EYFP would be expected to be greatly diluted immediately upon fusion, and might take time to re-accumulate to a high level.

In conclusion, our initial analysis of Musclebow fish provides robust data confirming the presence of numerous mpcs in each somite that proliferate on the surface of the myotome and constitute a stem cell pool that accounts for a significant fraction of the nuclei, and likely a majority of the non-fibre nuclei, in the larval somite.

### Mncs fuse to most fast fibres between 3 and 7 dpf

Musclebow allowed us to analyse the growth of the somitic myotome. The myotome grows by both increase in muscle fibre number (hyperplasia) and in muscle fibre size (hypertrophy) (Roy et al., 2017). However, the numerical increase in nuclei within fibres, and thus the total contribution of mpc terminal differentiation to growth has not been determined. Musclebow fish had marked mnc clusters in specific regions of particular somites (Fig. 4A). Detailed analysis of one such cluster showed not only increase in mnc number, but also terminal differentiation of mncs into fibres (Fig. 4B). Both fusion into larger pre-existing unmarked fast fibres and elongation of marked mncs into nascent fibres were observed within single clusters (Fig. 4B, blue and white arrowheads, respectively). Nascent fibres were defined by their thin and irregular transverse profile, extension for at least 70% somite length and were often located near the hypaxial or epaxial myotomal extremes (Fig. 3D). Quantification revealed up to fourfold increase in marked fast fibres between 3 and 6 dpf (Fig. 3C). In somite 14, for example, that had ten marked mncs, seven fast fibres (two mCer^+^ in epaxial myotome and five EYFP^+^ of which four were hypaxial and one epaxial) and three nascent fibres (all EYFP^+^ and in the hypaxial region) were marked at 3 dpf. By 6 dpf, there were 31 marked fibres in this somite, four mCer^+^ in each of the deep epaxial and hypaxial regions, six EYFP^+^ in the superficial epaxial domain and 17 EYFP^+^ in the hypaxial domain. One fast fibre in the superficial hypaxial domain was mCER^+^EYFP^+^, indicating fusion between myocytes derived from distinct mnc clones. It seems likely that the three hypaxial nascent EYFP^+^ fibres at 3 dpf matured into fast EYFP^+^ fibres at 6 dpf. Thus, at least ten additional EYFP^+^ mncs appear to have contributed to the ten additional hypaxial EYFP^+^ fast fibres, and five EYFP^+^ fibres were added in the epaxial domain. In addition to this contribution to pre-existing unmarked fibres, six superficial nascent EYFP^+^ fibres arose, one in the epaxial and five in the hypaxial domain. Strikingly, at 3 dpf, six and four EYFP^+^ mncs existed in the epaxial and hypaxial domains of this somite and this had increased to nine and 15, respectively, by 6 dpf, correlating with the abundant increase in EYFP^+^ fibres (Fig. 3C). Given the low frequency of labelling of mncs at 3 dpf and the significant increase in marked fast fibres, to about a third of all fibres in some somites by 6 dpf, we conclude that most fibres receive extra nuclei derived from mncs during this period.

### Stochastic clonal drift is common

Across five tracked somites, the average increase in marked fast fibres was 244% and in marked nascent fibres was 663%, which contrasted with a meagre 15% increase in marked slow fibres (Fig. 3C). Increase in marked fibres from 3 to 6 dpf generally correlated with the number of marked mncs present at 3 dpf (Figs 3C, S2A). Increases in marked fibres were accompanied by a 70% increase in marked mncs (Fig. 3C), indicating that mncs were proliferating at a sufficient rate to both expand their population and contribute many terminally-differentiated myocytes to the adjacent myotome.

Although, on average, mnc numbers increase, in somite 11 we observed a reduction from 14 to four marked mncs between 3 and 6 dpf, even as marked fast fibres increased from 19 to 46 and nascent fibres from three to 14 (Fig. 3C). This observation suggests that significant clonal drift can occur when proliferation of marked mncs fails to keep pace with their terminal differentiation. Indeed, comparison of the behaviour of the epaxial and hypaxial domains of somite 11 make this point clearly. Among EYFP^+^ mncs, 11 were epaxial and three hypaxial at 3 dpf. By 6 dpf, however, no EYFP^+^ mncs were detected in the epaxial domain, whereas four were still present in the hypaxial domain. The loss of 11 mncs from the epaxial domain was accompanied by an increase in EFYP^+^ fibres (both mature and nascent) of 23, from four to 27. In contrast, in the hypaxial domain, 15 additional EYFP^+^ fibres (from eight to 23) arose from the three original EYFP^+^ mncs, which themselves increased to four. Thus, in the hypaxial domain 16 nett EYFP^+^ cells were added, whereas in the epaxial domain, although there was a similar increase of 12 EYFP^+^ cells nett, it was at the expense of the entire epaxial mnc population. Complete loss of marked mncs was not observed in other somite halves (Fig. 3C). Thus, our data show how stochastic clonal drift can rapidly alter cell lineage when clone sizes are small.

### Uniform behaviour of mncs

Regional analysis of the fate of marked mncs between 3 and 6 dpf revealed several trends. First, there was a clear positive correlation (r = 0.77) between the number of marked mncs at 3 dpf and the number of newly-marked fibres accumulated between 3 and 6 dpf, which was discerned in both epaxial and hypaxial somite halves (Fig. S2A). Secondly, the presence of more marked mncs at 3 dpf showed a mild negative correlation (r = -0.39) with the number of extra mncs observed at 6 dpf (Fig. S2A), suggesting that regions with more mncs underwent less subsequent mnc proliferation. However, there was no correlation between the number of additional marked fibres and the number of extra marked mncs (r = -0.22), suggesting that extra differentiation did not account for the reduction in mncs. These trends were apparent equally in epaxial and hypaxial domains.

At 3 dpf, most marked mncs (86%) were located on the surface of the dermomyotome. By 6 dpf, in contrast, although marked mncs had more than doubled in number, most (59%) were at medial or deep levels within the myotome, consistent with the mpc migration previously described (Roy et al., 2017). The increase in marked fibres at different depths within the myotome was similar and correlated with the number of mncs in each somitic region at 3 dpf (Fig. S2B). In contrast, the increase in marked mncs at different depths was uncorrelated with the number of initial marked mncs. We conclude that apparently stochastic mnc proliferation and differentiation accompanied by invasion of the myotome leads to an increase in marked fibres and mncs and contributes to myotome growth over the early post-hatching period.

### Clones contribute to large regions of adult myotome

As some clones become large, their loss by stochastic neutral clonal drift is predicted to be less common (Klein and Simons, 2011). Congruently, live imaging of muscle clones in larval and adult animals revealed remarkable persistence and stability of clonal patterns. In a cohort of Musclebow-marked fish, blocks of muscle in individual somites were marked (Fig. 5A). Each fish showed a distinct pattern; marking was more common in trunk than tail, perhaps reflecting an increased number of mpcs in the larger trunk somites. No concordance was observed between somites on left and right sides of the animals. Marked muscle blocks ranged from a few tens of fibres to many hundred (Fig. 5A). As the *BB3mus7* line was made by Tol2-mediated insertion, it is anticipated to contain only a single copy of the BB3 transgene in the locus. Consistent with this, we find two primary colours of marked cell cluster: yellow (EYFP) or blue (mCer) (Fig. 5B). On rare occasions yellow and blue clusters partially overlap, showing that more than one early mpc contributes significantly to muscle growth in all regions observed (Fig. 5B,C). Although EYFP-marked regions were more numerous than mCer-marked regions, they displayed similar patch/clone sizes (Fig. 5B,C). This observation is consistent with the biased recombination in favour of EYFP observed in larvae.

**Fig. 5.**
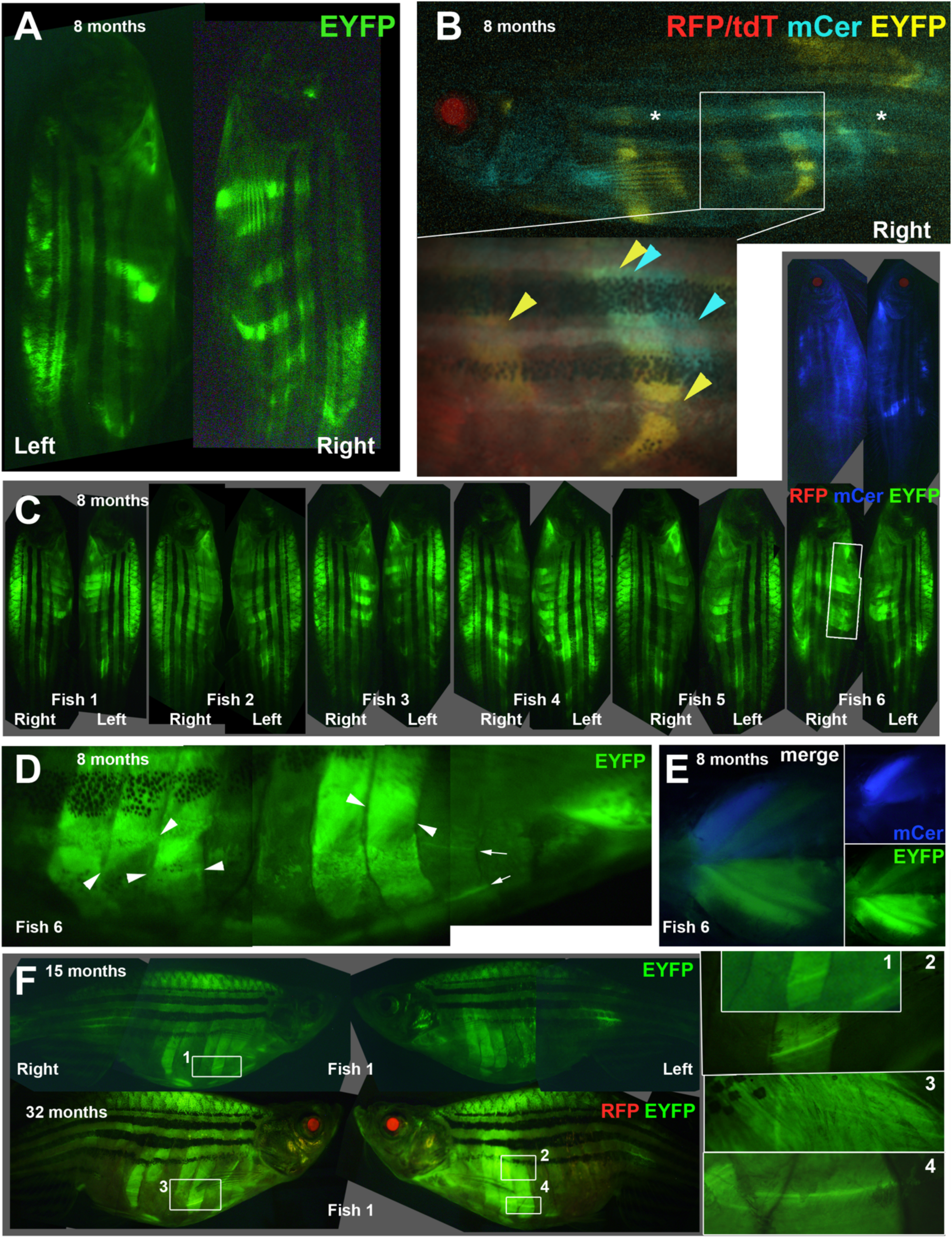
Stable clonal growth of muscle perdures throughout life. **A,B.** Single 8 month adult Musclebow-marked fish that were heat-shocked for 5 min at 37°C at 30% epiboly show distinct patterns of EYFP-marked myotomes on left and right side and between epaxial and hypaxial somite (A). When the right side of the same fish was imaged for all four fluorescent proteins, this fish had faint tdT throughout muscle and RFP in lens, indicating Cre expression, that had triggered clonal marking of several half somites in either EYFP or mCer. Asterisks indicate background blue autofluorescence from xanthophores. Box, magnified in inset, shows regions of hypaxial myotomes 11 and 14 marked by EYFP, whereas a region of somite 15 near the HZM is marked by mCer only. In a similar region of myotome 14, muscle is marked by both mCer and EYFP. **C-E.** Eight month old adult Musclebow-marked *BB3mus7* siblings (C) that were heatshocked for 5 min at 39°C at 24 hpf. Note the similar size but lower frequency of mCer clones compared to EYFP clones in Fish 6. Boxed region, magnified in D, reveals layers of fibres with distinct orientations (fibre ends indicated by pairs of arrowheads) and individual marked fibres (arrows). Ventral view of pectoral fin region of Fish 6 (E). **F.** Fish 1 from C at 15 and 32 months of age showing similar clonal pattern. Numbered boxes are magnified at right. Bar = 50 μm.

Where EYFP and mCer clones overlapped, some fibres contained both mCer and EYFP, whereas other single neighbouring fibres were uniquely marked, indicating that distinct clones gave rise to progeny that fused, as observed earlier in development (Figs 3 and 5B,C). Individual marked fibres within a myotome differed in EYFP intensity, consistent with the view that each region contained a mixture of marked and unmarked myoblasts whose fusion generated cells with differing levels of marker (Fig. 5D). These findings confirm that multinucleate fibres are generally formed by fusion of myoblasts from distinct clones.

Although hypaxial flank muscle was more readily imaged in adult fish than either epaxial or appendicular muscle, we observed no consistent difference in mpc behaviour in any muscle. Marked patches of both epaxial and, separately, hypaxial muscle were restricted to single myotomes/myotome domains, indicating that clones arise within single somites and their progeny do not significantly traverse the vertical myosepta throughout life (Fig. 5C). Similar regionalisation of clonal marking was observed in appendicular muscle (Fig. 5E).

Fibres both deep and superficial within the somite were marked in many clones, as reflected by their plane of focus and orientation to the body axis (Fig. 5D). Repeated imaging revealed that, once established during embryonic and larval growth, the marked regions did not notably change size (Fig. 5C,F). Nor did new labelled clones arise, suggesting that a) new recombination events due to leakage of the *HS:Cre* transgene are rare or do not occur during adult life and b) that either cell turnover was minimal or that the cells involved in muscle repair remained close to their clonally-related fibre. Moreover, individual fibres that were particularly bright, and therefore stood out from the group, were observed to persist for years (Fig. 5F). Thus, if there is significant on-going muscle repair during life in our aquarium, the clones that originally gave rise to normal muscle growth also contributed to muscle repair.

### Clonal analysis in muscle regeneration

To probe mpc behaviour during muscle regeneration, needle lesions were made in somitic muscle of 3-4 dpf *BB3mus7;HS:Cre.cryaa:RFP* fish in epaxial somite adjacent to regions containing clusters of EYFP-marked mncs within the body of the myotome (Fig. 6A). The clusters were likely clones because adjacent somites lacked EYFP mncs. For example, the animal shown in Fig. 6B-G had marked mnc clusters in just two of the 11 half somites scanned prior to wounding, which was representative of the whole larva. As the clusters had seven and eight mncs, respectively, the chance of this distribution arising without clonal relationship is approximately *p* = (2/11)^13^ = 2 x 10^-10^. Further, the probability that each cluster is a single clone is *p* ≥ 0.74 (the chance that, if there were even three marked clones, they would each be in separate somites). We can therefore examine the contribution of mnc clones arising during development to wound repair.

**Fig. 6.**
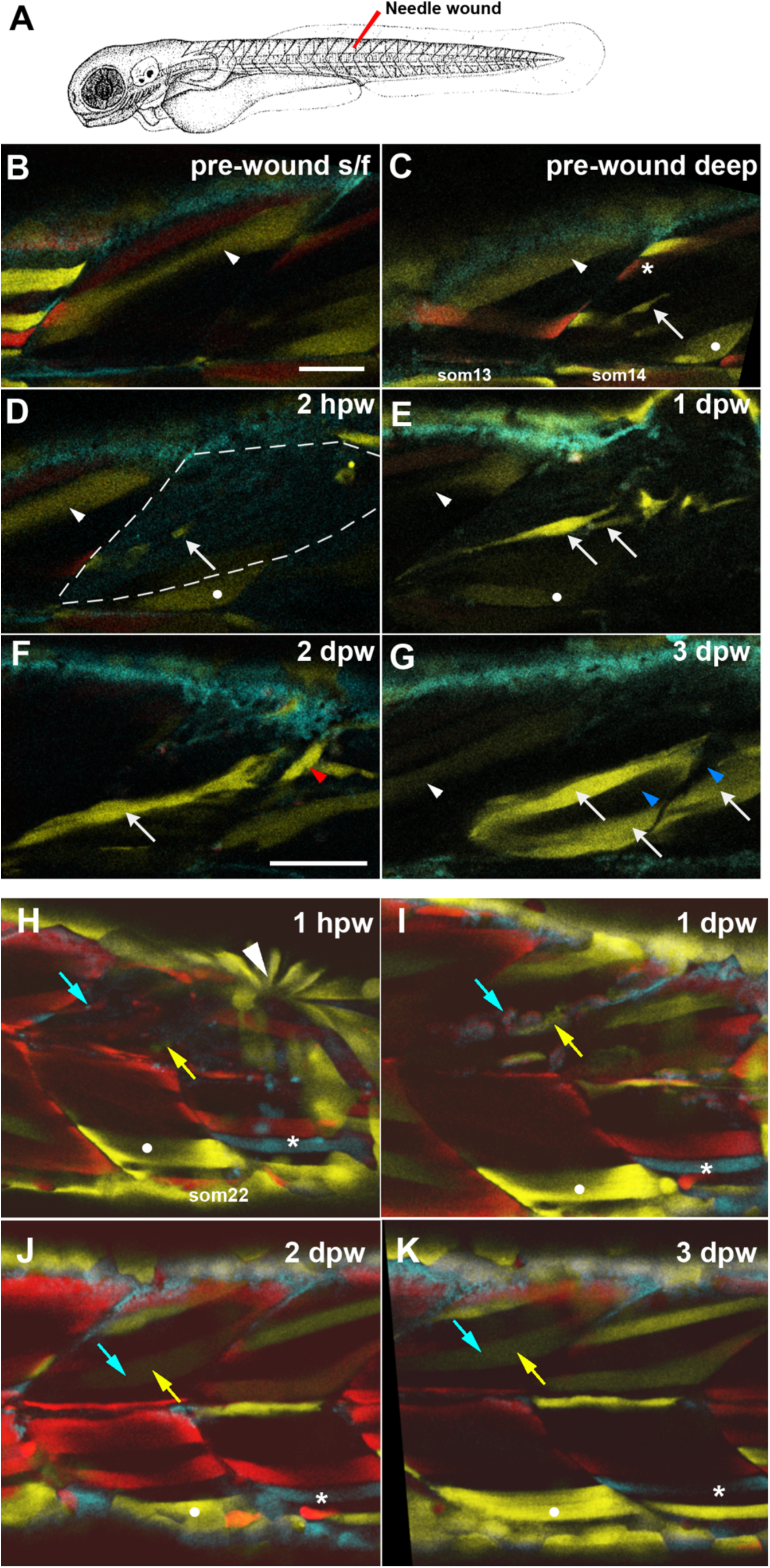
Partial contribution of clones to wound repair in zebrafish muscle. **A.** Schematic of wounding procedure (modified from (Kimmel et al., 1995)). A small needle wound was made near to marked mnc in somite 14 (som14) in the epaxial region of two adjacent somites at 4 dpf (B-G) or epaxial region of som16 at 3 dpf (H-K). Anterior to left, dorsal up. For clarity, short confocal stacks including the marked mnc are shown (B-G) extracted from full somite stacks before (B,C) and after (D-G, H-K) wounding. **B,C.** Superficial (B) and deeper (C) images before wounding. Distinct fibres present different colours after limited recombination at an early stage, thereby allowing unambiguous identification of the region (white arrowhead, white dot). A single EYFP-marked mpc (arrow) in som14 identified shortly prior to wounding. Note the presence of red (asterisk) and bright (arrowhead) and dim (dot) yellow fibres. **D.** By 2 h post-wounding (hpw), and despite dimming caused by the wounding procedure, the marked mnc (arrow) and the yellow fibres (arrowhead and dot) perdure in the wounded region (outlined by dashed line). In contrast, the red and bright yellow fibres have been lost. **E.** At 1 day post-wound (dpw), fluorescent signal has recovered and several bright EYFP-marked cells are detected in the wounded region (arrows). **F.** By 2 dpw, EYFP-marked cells are larger and elongating parallel to fibres (arrow) and present in adjacent unwounded somite (red arrowhead). **G.** At 3 dpw, the clusters of EYFP-marked mncs are replaced by several EYPF-marked fibres (arrows), indicating the extent of clonal contribution to the regenerate. Note the limited clonal expansion of the EYFP-marked clone over the three day period and the absence of label between the marked fibers (blue arrowheads). **H-K.** A second example showing single optical slices, with landmarks identifying wounded somites (white dot and asterisk) and tracking repair of a single fibre by fusion of cells from EYFP-marked (yellow arrows) and mCer/marked (cyan arrows) mpcs. Rapid purse-string epidermal wound closure is visible 1 hpw (arrowhead). Bars = 50 μm.

Comparison of confocal stacks immediately before and after wounding showed loss of some marked fibres (Fig. 6C asterisk), but persistence of other fibres (Fig. 6C-E white dot and arrowhead) and adjacent marked mncs (Fig. 6C-F white arrows). The same somite was then imaged daily during wound repair, and a number of striking cell behaviours were observed. First, marked mncs near the lesion contributed to regenerating fibres (Fig. 6E-G white arrows). Second, numbers of marked mncs did not detectably increase. Instead, mncs present on the day of wounding were often replaced by marked nascent fibres at 1 dpw (Fig. 6D,E white arrow). Such fibres matured and increased in size over subsequent days (Fig. 6F,G white arrows). These observations suggest that most marked mncs are mpcs and contribute rapidly to repair of damaged muscle.

We next asked whether regeneration occurred from rare stem cells within the somite, or was derived from a wider range of mpcs present prior to lesion. Regions of unlabelled fibres present within the regenerating domain suggested that distinct unlabelled mncs regenerated such fibres (Fig. 6G, blue arrowheads). Across a series of seven somites containing marked mncs, we observed appearance of marked fibres in the regenerated region in all cases (Table 3). On some occasions the marked mncs contributed to only a few fibres, but on other occasions the marked mncs gave rise to an increasingly large region of strongly EYFP^+^ regenerated fibres (Fig. 6G; Movie S2). Occasionally, newly-labelled fibres were observed in adjacent somites (Fig. 6F red arrowhead; Movie S2). In general, there was a positive correlation between the number of marked mncs prior to wounding and the extent of marked regenerated tissues (Table 3). In a second example, a single large regenerated fibre received contribution from two separate clones, respectively marked with mCer and EYFP (Fig. 6H-K). These findings show that multiple mpcs present within a single somite at the time of wounding contribute to wound repair.

**Table 3.** Contribution of marked mncs to wound repair.

| Stage at wounding (dpf<sup>#</sup>) | Wound location (somite number) | Marked mncs prior to wounding (number<sup>§</sup>) | Regeneration analysed (dpw) | Marked regenerated fibers (number*) |
| --- | --- | --- | --- | --- |
| 3 | 6 | Unrecorded | 4 | 2 or 3 |
| 3 | 19 | 2 to 3 | 3 | 2 |
| 3 | 22 | 3 to 4 | 1 | 4 |
|  |  |  | 2 | + 2 additional fibres |
|  |  |  | 3 | + 2 extra thin fibres |
|  |  |  | 3 | = 8 in fibres total |
| 3 | 13-16 | 2 to 3 | 3 | 2 to 3 |
| 3 | 18-20 | 3 to 4 | 1 | 2 |
| 4 | 15 (+14 partially) | > 4 | 3 | 2 to 3 |
| 4 | 14 | 2 to 3 | 1 | 3 to 4 |
|  |  |  | 2 | bigger and brighter |
|  |  |  |  | + 4 additional fibres |
|  |  |  | 3 | bigger and brighter |
|  |  |  |  | + at least 5 additional fibres |
|  |  |  | 3 | = 12 or 13 fibres total |

<sup>#</sup> Seven fish from separate lays were wounded near marked mncs on different dates over a 5 month period. The wounded region was tracked over subsequent days by 4D confocal microscopy.
<sup>§</sup> When mncs touch it is not possible to be sure of the number.
\* When marked fibres align parallel and adjacent it was sometimes hard to be sure of their number.

## Discussion

We have investigated clonal cell behaviour during both the growth and repair of zebrafish muscle in the initial phases of myogenesis and in adult life. Our findings reveal that: 1. A cohort of early non-proliferative mpcs generate the early myotome. 2. Marked mpc clones contribute little to muscle growth prior to hatching. 3. At 3 dpf when larvae have hatched, around 25% the nuclei in a single somite are in mncs. 4. From 3-6 dpf, dermomyotome-derived mpc clones rapidly expand while some progeny undergo terminal differentiation. 5. Neither fibre nor mpc death was observed in uninjured animals. 6. Individual marked muscle fibres persist across much of the lifespan indicating low rates of nuclear turnover. 7. In adulthood, early-marked clones label stable blocks of tissue comprising a significant fraction of either epaxial or hypaxial somite, in one or a few adjacent somites. 8. Fusion of cells from separate early-marked clones occurs in regions of clone overlap. 9. Wounds made next to marked mpc clones are regenerated partially from the marked cells, suggesting that many/most mpcs (both marked and unmarked) within a muscle block contribute to local wound repair.

### Variation in clonal persistence during embryonic myogenesis

Our analysis revealed no clustering of labelled early embryonic myofibres with similarly-labelled mnc clones. This finding confirms the observations from dye-filling experiments that most early somitic cells differentiate directly into either of two early populations of slow and fast embryonic myofibres with little or no proliferation prior to 2 dpf (Devoto et al., 1996; Hirsinger et al., 2004; Hollway et al., 2007; Stellabotte et al., 2007). We nevertheless find that a small subpopulation of early-marked somitic cells persist as mncs, presumably derived from somite anterior border cells (Devoto et al., 2006; Hammond et al., 2007; Hollway et al., 2007; Stellabotte et al., 2007).

These mncs, numbering at least 15/somite, subsequently proliferate rapidly and generate mpc clones within which individual cells undergo terminal differentiation into muscle fibres after hatching. Other cells in the same marked mpc clones remain as proliferative mncs at least until 13 dpf, the longest timelapse we have performed. Many such marked-mnc clusters are initially located superficially within the somite in the region described variously as dermomyotome or external cell layer (Devoto et al., 2006).

We did not observe marked mnc clusters generating both muscle fibres and another obvious distinct non-myotomal cell type, such as sclerotome or dermis. Although the *BB3mus7* Musclebow transgene does not express in non-myogenic tissue, one would expect perdurance of EYPF and mCer marker proteins for some days, as we have seen with a *pax7a*-driven GFP transgene (Pipalia et al., 2016). We therefore hypothesise that most early dermomyotomal cells have the capacity to make proliferative mpcs but do not generate abundant non-myogenic types, at least in the midbody regions and larval stages we have mainly investigated.

Our analysis of the cellular content of the zebrafish larval somite begins a quantitative understanding of proliferation, differentiation and growth of cells within this defined tissue block. Change in number of som16 mncs from around 40 Pax3/7+ve mpcs at 24 hpf (Hammond et al., 2007), to about 90 mncs at 72 hpf suggests proliferation. Indeed, proliferation was observed in marked mnc clones. We previously showed that each som12-16 has around 271±19 nuclei at 24 hpf, and that both Pax3/7^+^ and Pax3/7^-^ cells within the somite proliferate as indicated by phospho-Histone H3 and EdU staining (Hammond et al., 2007; Roy et al., 2017). Given that som12 is larger than som16 throughout development, and that our current data indicate that som16 has 301 nuclei at 3 dpf, the data suggest that mncs have proliferated and the nuclei generated have been retained within the somite. Indeed, based on Myogenin co-expression, we have previously shown that Pax3/7+ve mncs are undergoing differentiation into fibres at 3 dpf and beyond (Roy et al., 2017). Moreover, proliferation of mnc clones and increase in mnc numbers occurs between 1 and 3 dpf. Thus, an increase in mncs, despite some differentiation into fibres is likely. Making the modest assumptions that i) som16 had 240 cells at 24 hpf rising to 300 at 3 dpf, ii) most proliferating cells within the somite are mpcs and iii) that all new fibre nuclei arise from mpcs within the somite, calculation suggests that on average 60% of mpcs divide each day whereas just 6.25% differentiate each day in embryonic somites.

Marked-mpc clones show heterogenous behaviour in larval life. Some marked-mpc clones contribute to small presumed-nascent fibres, particularly at the dorsal and ventral somitic extremes after 3 dpf. Other marked mpc clones given rise to cells that invade the myotome after about 3 dpf and then contribute to pre-existing fast fibres through cell fusion. Many marked mncs show little division, whereas some proliferate well above average. Although, we have not yet analysed sufficient clones in detail, our data are consistent with the view that the decision between proliferation, migration and terminal differentiation within mpc clones is indeterminate and stochastic. By this we mean that either it is a truly probabilistic process intrinsic to the mpc, or that acute local cues influence the decision, for example signals from nearby fibres.

### Cell death is rare during muscle development

In amniotes, cell death is often employed to control cell numbers, drive morphogenesis, and correct errors in developmental progression. For example, approximately twice the required final number of motoneurons are generated, with half dying after the initial innervation of target muscle fibres ((Buss et al., 2006) and references therein). During the course of our studies some thousands of individually-marked and identifiable muscle fibres and mncs were examined. No instances of marked-cell disappearance, fibre thinning/atrophy, fibre detachment from somite borders or phagocytosis were observed, suggesting that cell death within early myogenic lineages is not generally employed to sculpt myogenic progression in zebrafish.

### No apparent cell lineage basis for early myogenesis

Studies over fifty years have suggested that myoblast intrinsic heterogeneity could underlie formation of distinct muscle fibre types and/or generations (Cossu et al., 1988; Hauschka, 1974; Hughes and Salinas, 1999; Miller and Stockdale, 1986). One motivation for the development of Musclebow was the hope of revealing diverse populations of mpcs showing distinct categories of behaviour during the early phases of myogenesis, which have been hard to examine in amniotes. No such diversity has been apparent to date; while diverse, clones differ in behaviour over a smooth spectrum in all respects considered. We note, however, that the number of clones examined and the duration of time-lapse observations in our initial study is limited. Therefore, divergent mpc lineages present at low abundance or in locations difficult to image, such as dorsal and ventral somitic extremes, may not have been sufficiently sampled. Moreover, almost all amniote studies have been performed on limb muscle, whereas we have focused on intrinsic somitic muscle, which is homologous to the deep muscle of the back.

### Clonal drift and persistence in adult life

Time-lapse imaging in early larval development reveals variation in mpc clonal behaviour, such that some clones become large, while others fail to endure as mpcs. We find no reason not to interpret this observation as the result of stochastic clonal drift. After larval growth and into the adult stage, such large mpc clones give rise to patches of marked muscle fibres extending over significant portions of the epaxial or hypaxial myotome. As adult zebrafish muscle fibres have hundreds of nuclei (Ganassi et al., 2020), it is likely that small numbers of fusing nuclei from a marked mpc clone can label essentially all fibres in a region. Regions of overlap between fibres labelled with both EYFP and mCer prove that adult fibres are of multiclonal origin.

Adjacent somites on the anteroposterior axis, somites at the same anteroposterior position on left and right sides, and epaxial and hypaxial domains of single somites appear not to share large mpc clones. Labelling of adjacent regions is what might be expected by the chance of coincident recombination, suggesting that mpcs rarely cross the vertical or horizontal myosepta during normal life.

Once a large region of a myotome is labelled, that labelling persists throughout life and does not appear to spread. In principle, such observations could reflect either low turnover of muscle nuclei or that once marked mpcs have become abundant they perdure throughout life and contribute to ongoing regeneration in the same region that they originally marked fibres. However, as wounding studies ((Gurevich et al., 2016) and see below) suggest that single mpc clones can make significant contribution to larval muscle wounds through clonal drift, our observations suggest that regeneration is not extensive in adult life. If such regeneration were occurring, a gradual expansion of at least some marked regions would be expected, which was not observed. Moreover, single strongly-marked fibres in locations such as the ventral body wall favourable for adult imaging, were observed to endure for years. These observations strongly argue that turnover in adult zebrafish muscle is not extensive, as had been described in humans (Spalding et al., 2005).

### Multiple nearby mpcs contribute to muscle regeneration

Wounds made next to marked mpcs are generally regenerated with a partial contribution of the marked cell to the repaired fibres, so long as the marked mpc survived the wounding procedure. Given the low rate of mpc marking by Musclebow, our data argue against the existence in larval life of a minor population of stem mpcs responsible for most fibre regeneration, as has previously been suggested (Gurevich et al., 2016). As such stem cells would have been unmarked in most wounded somites, the regenerated fibres would rarely be marked; this was not the case. Moreover, we observe regeneration of wounds in which only a fraction of fibres within the regenerated region are labelled, again arguing for polyclonal mpc regeneration and against a unique regenerative stem cell. How, then, could the clear-cut appearance of monoclonal regeneration (Gurevich et al., 2016) have arisen? We suggest that i) widespread labelling of mpcs from *msgn1:CreERT2* (which is only expressed in early presomitic mesoderm (http://zfin.org/ZDB-GENE-030722-1/expression), ii) recombination bias towards EYFP in *ubi:Zebrabow* similar to what we observe in *BB3mus7* Musclebow (both Zebrabow and Musclebow are based on the same *Brainbow-1.0 ‘L’* vector (Pan et al., 2013)) and iii) clonal drift during early life led to the erroneous conclusion of clonal regeneration (Gurevich et al., 2016). Similarly, regeneration in adult mammalian muscle is clearly polyclonal (Tierney et al., 2018).

### Novel Musclebow technology

By using a heat-shock promoter to trigger recombination in individual cells irrespective of their intrinsic gene expression, our approach differs from those employing cell type-restricted Cre expression (Gurevich et al., 2016; Nguyen and Currie, 2018; Nguyen et al., 2017). Instead, our cell labelling is driven by insertion of an enhancer trap vector into a genomic locus. Reverse PCR with Tol2 primers has suggested that the *BB3mus7* transgene is inserted into a common genomic repeat element that has so far precluded unambiguous mapping. [XXGenetic mapping reveals that in our line the BB3 transgene insertion is located in an intron of the *brsk2a* gene on chromosome 25 that may have undergone some rearrangement. As expression of *brsk2a* is not normally restricted to myogenic tissue, it is possible that the reorganized locus may have facilitated an interaction of a normally distant muscle enhancer with the BB3 basal promoter.] In addition, a second copy of the BB3 transgene may be present in a linked locus and show widespread low level expression. Nevertheless, as we have recently been able to separate the two loci by extensive outcrossing, we think additional copies do not contribute to muscle labelling significantly. The *BB3mus7* line may prove useful in a variety of situations, such as transplantations or lineage tracing studies, where marking of mpcs and fibres is required.

## Acknowledgements

We thank Hughes lab members, Esperanza Hughes-Salinas, and the staff of the NIMR, Francis Crick Institute and KCL fish facilities.

## Supplementary Data

**Fig. S1. Generation of the *kg330* allele.** Linked BB3 insertions in *kg309* yield distinct patterns of recombination. When four independent female *BB3mus7^kg309/+^*;*HS:Cre.cryaa:RFP^kg310^* fish were outcrossed to wild type fish, only 182/571 (32%) of progeny lacked fluorescence (excluding lens RFP), a highly significant difference from both the predicted 50% for a single transgene insertion (p = 9.0E-10, *X*^2^ test) and the 25% expected for two (or <25% for more than two) separate unlinked transgenes (p = 1.5E-4, *X*^2^ test) indicating two linked BB3 insertions in the *BB3mus7^kg309^* allele. Female germline Cre expression leads to non-mosaic recombination yielding several distinct patterns of fluorescent protein expression at each linked BB3 locus. **A**. Recombination of the *BB3mus7* transgene yielded strong myotomal EYFP expression (arrow). A second separable insertion yielded widespread low-level EYFP (*BB3weak*) expression in most tissues including brain, eye, heart, epidermis and gut (arrowheads). Such fish never express mCer (lower panel). **B.** Many fish lacked fluorescence altogether, revealing the background autofluorescence level. **C,D.** Less frequently, individuals recombined at *BB3mus7* to give myotomal mCer expression (arrows) either alone (C lower fish and D) or in combination with *BB3weak* EYFP (C upper fish). Note the presence of *BB3weak* signal (filled arrowheads) in head of the upper individual, but not of the lower individual (open arrowheads) and *BB3mus7* mCer in both upper (arrow) and lower (blue arrows) larvae. **E.** Individuals with only the *BB3weak* EFYP were numerous. Note the contrast of the weak signal in trunk in this enhanced imaged compared with the *BB3mus7EYFP,BB3weakEYFP* transgenic in panel A. **F.** Frequency of each genotype indicating genetic linkage and recombination dynamics. Note that *BB3mus7* transmitted to 274/571 (48%) of progeny, as expected for a single heterozygous Mendelian locus (p = 0.34, *X*^2^ test). Lack of mCer recombination at the *BB3weak* locus suggests genetic interference, supporting linkage of *BBSmus7* and *BB3weak.* **G.** Outcrossing of *kg309* (tdT expression shown in the upper individual) permitted separation on the basis of the tdTomato of *BB3mus7* and *BB3weak* to isolate the *BB3mus7only kg330* allele (lower larva). Note the widespread weak tdT signal in head and gut of *BB3mus7^kg309^* (arrowheads) and its reduction in *BB3mus7^kg330^* compared to the *BB3mus7* myotomal signal (arrows). **H.** Lateral views of single *BB3mus7^kg330^* and *HS:Cre.cryaa:RFP^kg310^* fish at 6 dpf showing the distinct RFP and tdT signal and the low mCer and EYFP background in somites of unrecombined Musclebow.

**Fig. S2. Regional correlation of myogenesis from marked mncs.** Change in number of marked fibres (upper graphs) and mncs (lower charts) in each region of six single somites analysed at both 3 and 6 dpf, from same dataset shown in Fig. 3C. **A.** Similar behaviour in epaxial and hypaxial somitic regions. **B.** Dispersion of largely superficial mncs into deeper regions. Colours indicate final location at 6 dpf.

**Movie S1. Stability of Musclebow labelling on short timescales.** *Tg(BB3mus7)^kg309^;Tg(HS:Cre.cryaa:RFP)^kg310^* embryo was subjected to 5 min heat-shock at 30% epiboly. After embedding in 3% methylcellulose with light tricaine at 26 hpf, 15 sequential confocal time-lapse 3-colour 3-tile 2 μm stacks were taken every 24 min over 6 h at 24°C. An 11-slice mip of each stack is shown.

**Movie S2. Rotation of healed wound.** *Tg(BB3mus7)^kg309^;Tg(HS:Cre.cryaa:RFP)^kg310^* embryo was subjected to 5 min heat-shock at 24 hpf, a selected region containing a marked mnc wounded at 4 dpf and allowed to regenerate for 3 days while repeatedly observed (also shown in Fig. 6B-F). A 2-colour confocal stack at 3 dpw (top) was segmented (bottom) to reveal the marked-mnc-derived regenerated fibres in som14.

## References

1. Blau, H. M., Pavlath, G. K., Hardeman, E. C., Chiu, C. P., Silberstein, L., Webster, S. G., Miller, S. C. and Webster, C. (1985). Plasticity of the differentiated state. Science 230, 758–766.

2. Buss, R. R., Sun, W. and Oppenheim, R. W. (2006). Adaptive roles of programmed cell death during nervous system development. Annu Rev Neurosci 29, 1–35.

3. Collins, C. A., Olsen, I., Zammit, P. S., Heslop, L., Petrie, A., Partridge, T. A. and Morgan, J. E. (2005). Stem cell function, self-renewal, and behavioral heterogeneity of cells from the adult muscle satellite cell niche. Cell 122, 289–301.

4. Cooper, M. S., Szeto, D. P., Sommers-Herivel, G., Topczewski, J., Solnica-Krezel, L., Kang, H. C., Johnson, I. and Kimelman, D. (2005). Visualizing morphogenesis in transgenic zebrafish embryos using BODIPY TR methyl ester dye as a vital counterstain for GFP. Dev Dyn 232, 359–368.

5. Cossu, G., Ranaldi, G., Senni, M. I., Molinaro, M. and Vivarelli, E. (1988). ’Early’ mammalian myoblasts are resistant to phorbol ester-induced block of differentiation. Development 102, 65–69.

6. Devoto, S. H., Melancon, E., Eisen, J. S. and Westerfield, M. (1996). Identification of separate slow and fast muscle precursor cells in vivo, prior to somite formation. Development 122, 3371–3380.

7. Devoto, S. H., Stoiber, W., Hammond, C. L., Steinbacher, P., Haslett, J. R., Barresi, M. J., Patterson, S. E., Adiarte, E. G. and Hughes, S. M. (2006). Generality of vertebrate developmental patterns: evidence for a dermomyotome in fish. Evol Dev 8, 101–110.

8. DiMario, J. X., Fernyak, S. E. and Stockdale, F. E. (1993). Myoblasts transferred to the limbs of embryos are committed to specific fibre fates. Nature 362, 165–167.

9. DiMario, J. X. and Stockdale, F. E. (1995). Differences in the developmental fate of cultured and noncultured myoblasts when transplanted into embryonic limbs. Experimental Cell Research 216, 431–442.

10. Ganassi, M., Badodi, S., Wanders, K., Zammit, P. S. and Hughes, S. M. (2020). Myogenin is an essential regulator of adult myofibre growth and muscle stem cell homeostasis. Elife 9, e60445.

11. Gearhart, J. D. and Mintz, B. (1972). Clonal origins of somites and their muscle derivatives: evidence from allophenic mice. Dev Biol 29, 27–37.

12. Gros, J., Manceau, M., Thome, V. and Marcelle, C. (2005). A common somitic origin for embryonic muscle progenitors and satellite cells. Nature 435, 954–958.

13. Gurevich, D. B., Nguyen, P. D., Siegel, A. L., Ehrlich, O. V., Sonntag, C., Phan, J. M., Berger, S., Ratnayake, D., Hersey, L., Berger, J., et al. (2016). Asymmetric division of clonal muscle stem cells coordinates muscle regeneration in vivo. Science 353, aad9969.

14. Haldar, M., Karan, G., Tvrdik, P. and Capecchi, M. R. (2008). Two Cell Lineages, myf5-Dependent and myf5-Independent, Participate in Mouse Skeletal Myogenesis. Dev Cell 14, 437-445.

15. Hammond, C. L., Hinits, Y., Osborn, D. P., Minchin, J. E., Tettamanti, G. and Hughes, S. M. (2007). Signals and myogenic regulatory factors restrict pax3 and pax7 expression to dermomyotome-like tissue in zebrafish. Dev Biol 302, 504–521.

16. Hans, S., Freudenreich, D., Geffarth, M., Kaslin, J., Machate, A. and Brand, M. (2011). Generation of a non-leaky heat shock-inducible Cre line for conditional Cre/lox strategies in zebrafish. Dev Dyn 240, 108–115.

17. Hauschka, S. D. (1974). Clonal analysis of vertebrate myogenesis. 3. Developmental changes in the muscle-colony-forming cells of the human fetal limb. Dev Biol 37, 345-368.

18. Hirsinger, E., Stellabotte, F., Devoto, S. H. and Westerfield, M. (2004). Hedgehog signaling is required for commitment but not initial induction of slow muscle precursors. Dev Biol 275, 143–157.

19. Hollway, G. E., Bryson-Richardson, R. J., Berger, S., Cole, N. J., Hall, T. E. and Currie, P. D. (2007). Whole-somite rotation generates muscle progenitor cell compartments in the developing zebrafish embryo. Dev Cell 12, 207–219.

20. Hughes, S. M. (1999). Fetal myoblast clones contribute to both fast and slow fibres in developing rat muscle. Int J Dev Biol 43, 149–155.

21. Hughes, S. M. and Blau, H. M. (1990). Migration of myoblasts across basal lamina during skeletal muscle development. Nature 345, 350–353.

22. Hughes, S. M. and Blau, H. M. (1992). Muscle fiber pattern is independent of cell lineage in postnatal rodent development. Cell 68, 659–671.

23. Hughes, S. M. and Salinas, P. C. (1999). Control of muscle fibre and motoneuron diversification. Curr Opin Neurobiol 9, 54–64.

24. Hutcheson, D. A., Zhao, J., Merrell, A., Haldar, M. and Kardon, G. (2009). Embryonic and fetal limb myogenic cells are derived from developmentally distinct progenitors and have different requirements for beta-catenin. Genes Dev 23, 997–1013.

25. Kassar-Duchossoy, L., Giacone, E., Gayraud-Morel, B., Jory, A., Gomes, D. and Tajbakhsh, S. (2005). Pax3/Pax7 mark a novel population of primitive myogenic cells during development. Genes Dev 19, 1426–1431.

26. Kelu, J. J., Pipalia, T. G. and Hughes, S. M. (2020). Circadian regulation of muscle growth independent of locomotor activity. Proc Natl Acad Sci U S A 117, 31208–31218.

27. Kimmel, C. B., Ballard, W. W., Kimmel, S. R., Ullmann, B. and Schilling, T. F. (1995). Stages of embryonic development of the zebrafish. Developmental Dynamics 203, 253–310.

28. Klein, A. M. and Simons, B. D. (2011). Universal patterns of stem cell fate in cycling adult tissues. Development 138, 3103–3111.

29. Livet, J., Weissman, T. A., Kang, H., Draft, R. W., Lu, J., Bennis, R. A., Sanes, J. R. and Lichtman, J. W. (2007). Transgenic strategies for combinatorial expression of fluorescent proteins in the nervous system. Nature 450, 56–62.

30. Mahalwar, P., Walderich, B., Singh, A. P. and Nüsslein-Volhard, C. (2014). Local reorganization of xanthophores fine-tunes and colors the striped pattern of zebrafish. Science 345, 1362–1364.

31. Messina, G., Biressi, S., Monteverde, S., Magli, A., Cassano, M., Perani, L., Roncaglia, E., Tagliafico, E., Starnes, L., Campbell, C. E., et al. (2010). Nfix Regulates Fetal-Specific Transcription in Developing Skeletal Muscle. Cell 140, 554–566.

32. Miller, J. B. and Stockdale, F. E. (1986). Developmental regulation of the multiple myogenic cell lineages of the avian embryo. J. Cell Biol. 103, 2199–2208.

33. Motohashi, N., Uezumi, A., Asakura, A., Ikemoto-Uezumi, M., Mori, S., Mizunoe, Y., Takashima, R., Miyagoe-Suzuki, Y., Takeda, S. and Shigemoto, K. (2019). Tbx1 regulates inherited metabolic and myogenic abilities of progenitor cells derived from slow- and fast-type muscle. Cell death and differentiation 26, 1024–1036.

34. Nguyen, P. D. and Currie, P. D. (2018). Guidelines and best practices in successfully using Zebrabow for lineage tracing multiple cells within tissues. Methods 150, 63–67.

35. Nguyen, P. D., Gurevich, D. B., Sonntag, C., Hersey, L., Alaei, S., Nim, H. T., Siegel, A., Hall, T. E., Rossello, F. J., Boyd, S. E., et al. (2017). Muscle Stem Cells Undergo Extensive Clonal Drift during Tissue Growth via Meox1-Mediated Induction of G2 Cell-Cycle Arrest. Cell stem cell 21, 107–119 e106.

36. Pan, Y. A., Freundlich, T., Weissman, T. A., Schoppik, D., Wang, X. C., Zimmerman, S., Ciruna, B., Sanes, J. R., Lichtman, J. W. and Schier, A. F. (2013). Zebrabow: multispectral cell labeling for cell tracing and lineage analysis in zebrafish. Development 140, 2835–2846.

37. Pipalia, T. G., Koth, J., Roy, S. D., Hammond, C. L., Kawakami, K. and Hughes, S. M. (2016). Cellular dynamics of regeneration reveals role of two distinct Pax7 stem cell populations in larval zebrafish muscle repair. Dis Model Mech 9, 671–684.

38. Quinn, L. S., Holtzer, H. and Nameroff, M. (1985). Generation of chick skeletal muscle cells in groups of 16 from stem cells. Nature 313, 692–694.

39. Robson, L. G. and Hughes, S. M. (1999). Local signals in the chick limb bud can override myoblast lineage commitment: induction of slow myosin heavy chain in fast myoblasts. Mech Dev 85, 59–71.

40. Roy, S. D., Williams, V. C., Pipalia, T. G., Li, K., Hammond, C. L., Knappe, S., Knight, R. D. and Hughes, S. M. (2017). Myotome adaptability confers developmental robustness to somitic myogenesis in response to fibre number alteration. Dev Biol 431, 321–335.

41. Schafer, D. A., Miller, J. B. and Stockdale, F. E. (1987). Cell diversification within the myogenic lineage: in vitro generation of two types of myoblasts from a single progenitor cell. Cell 48, 659–670.

42. Seger, C., Hargrave, M., Wang, X., Chai, R. J., Elworthy, S. and Ingham, P. W. (2011). Analysis of Pax7 expressing myogenic cells in zebrafish muscle development, injury, and models of disease. Dev Dyn 240, 2440–2451.

43. Shaner, N. C., Campbell, R. E., Steinbach, P. A., Giepmans, B. N., Palmer, A. E. and Tsien, R. Y. (2004). Improved monomeric red, orange and yellow fluorescent proteins derived from Discosoma sp. red fluorescent protein. Nat Biotechnol 22, 1567–1572.

44. Spalding, K. L., Bhardwaj, R. D., Buchholz, B. A., Druid, H. and Frisen, J. (2005). Retrospective birth dating of cells in humans. Cell 122, 133–143.

45. Stellabotte, F., Dobbs-McAuliffe, B., Fernandez, D. A., Feng, X. and Devoto, S. H. (2007). Dynamic somite cell rearrangements lead to distinct waves of myotome growth. Development 134, 1253–1257.

46. Stockdale, F. E. and Holtzer, H. E. (1961). DNA synthesis and myogenesis. Exp. Cell Res. 24, 508–520.

47. Tierney, M. T., Stec, M. J., Rulands, S., Simons, B. D. and Sacco, A. (2018). Muscle Stem Cells Exhibit Distinct Clonal Dynamics in Response to Tissue Repair and Homeostatic Aging. Cell stem cell 22, 119–127 e113.

48. Westerfield, M. (2000). The Zebrafish Book - A guide for the laboratory use of zebrafish (Danio rerio): University of Oregon Press.

49. Windner, S. E., Bird, N. C., Patterson, S. E., Doris, R. A. and Devoto, S. H. (2012). Fss/Tbx6 is required for central dermomyotome cell fate in zebrafish. Biology open 1, 806–814.

50. Cooper, M. S., Szeto, D. P., Sommers-Herivel, G., Topczewski, J., Solnica-Krezel, L., Kang, H. C., Johnson, I. and Kimelman, D. (2005). Visualizing morphogenesis in transgenic zebrafish embryos using BODIPY TR methyl ester dye as a vital counterstain for GFP. Dev Dyn 232, 359–368.

51. Kimmel, C. B., Ballard, W. W., Kimmel, S. R., Ullmann, B. and Schilling, T. F. (1995). Stages of embryonic development of the zebrafish. Developmental Dynamics 203, 253–310.

